# OpenBase: a universal framework for high-accuracy single-molecule detection of diverse non-canonical DNA bases using nanopore sequencing

**DOI:** 10.64898/2026.05.13.724850

**Authors:** Zhen-Dong Zhong, Ying-Yuan Xie, Ru-Jia Luo, Hong-Xuan Chen, Zhi-Shuo Qiao, Ze-Hui Ren, Zhang Zhang, Guan-Zheng Luo

**Affiliations:** State Key Laboratory of Biocontrol, MOE Key Laboratory of Gene Function and Regulation, Guangdong Province Key Laboratory of Pharmaceutical Functional Genes, School of Life Sciences, Sun Yat-sen University, Guangzhou 510275, China; Sun Yat-sen University Institute of Advanced Studies Hong Kong, Science Park, Hong Kong SAR, 999077, China; Innovation Center for Evolutionary Synthetic Biology, Sun Yat-sen University, Guangzhou 510275, China

## Abstract

Nanopore sequencing holds great potential for the direct detection of non-canonical DNA bases from electrical signals, yet current approaches remain limited to a few classical epigenetic marks. Here we present OpenBase, an open and universal framework that standardizes training-data generation and deep learning based-modeling for single-molecule identification of diverse non-canonical DNA bases. OpenBase transforms nanopore sequencing into a broadly accessible platform for exploring the chemical diversity of DNA.

## Main

Nanopore sequencing provides a direct, single-molecule view of DNA chemistry by recording electrical signals that reflect the identity and chemical state of each nucleotide^1-3^. This technology has the potential to detect a wide spectrum of non-canonical bases, thereby offering unprecedented access to the chemical and epigenetic information in DNA. Nevertheless, current approaches remain limited to a few classical marks, such as 5mC, 5hmC, and 6mA^4-6^, largely due to the lack of standardized training data and generalizable models. Beyond these classical marks, DNA harbors a wide variety of non-canonical bases, encompassing not only endogenous epigenetic modifications but also exogenous damage-derived bases and synthetic analogs that hold significant promise for applications in synthetic biology^7-12^. Yet, the broader potential of nanopore sequencing to comprehensively decode the DNA base spectrum remains largely unrealized.

To address this gap, we developed OpenBase, an open and universal framework for high-accuracy detection of diverse non-canonical DNA bases at the single-molecule level (Fig. 1a). The workflow comprises three core modules: (i) standardized training-data generation, (ii) nanopore signal acquisition, and (iii) deep learning-based model training. Training data are generated from *in vitro-*synthesized DNA oligonucleotides composed of a fixed 5’ region, a randomized central sequence, and a 3’ reverse-complementary region.

**Fig. 1:**
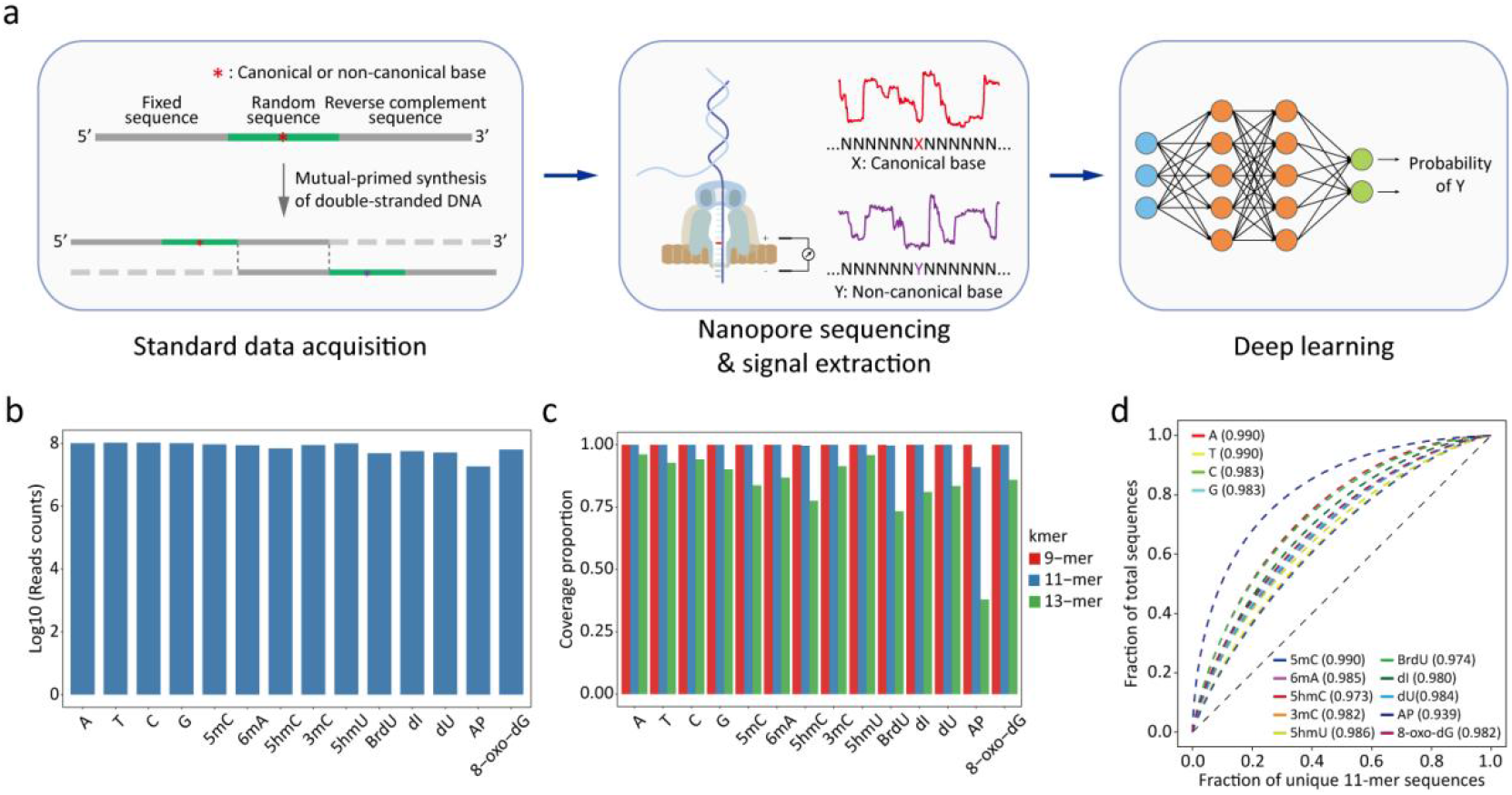
OpenBase workflow and dataset statistics. **a**, Schematic of the three core modules of OpenBase. **b-d**, Dataset characteristics across 14 base types, including reads counts (**b**), sequence-type coverage (**c**), and 11-mer uniformity (**d**, quantified by the evenness index).

The target base (canonical or non-canonical) is placed at the center of the random region, while the complementary 3’ tail serves as both primer and template during annealing and extension to form a double-stranded structure. The duplex products are then subjected to nanopore sequencing, from which raw current signals corresponding to each base type are extracted for model training. Each non-canonical base is paired with its canonical counterpart to train a base-classification model using a hybrid deep learning architecture.

By incorporating a reverse-complementary 3’ tail, OpenBase minimizes dependence on oligonucleotide synthesis length while fully meeting the requirements of nanopore sequencing. The designed single-stranded oligonucleotide is approximately 120 nt in length, yielding a ∼200 nt double-stranded product after primer extension (Extended Data Fig. 1a-b). The randomized central region, flanked by 16 random nucleotides on each side of the target base, ensures high sequence diversity. Both terminal regions (∼40 nt each) serve as robust alignment anchors and facilitate efficient duplex formation (Extended Data Fig. 2). This design enables large-scale generation of compositionally balanced datasets covering 14 distinct base types (including canonical bases), providing a solid foundation for high-quality model training (Fig. 1b-d and Extended Data Fig. 3).

For model development, 5mC and 6mA were selected as representative examples to construct and validate OpenBase. The model architecture combines convolutional neural networks (CNNs) with Transformer modules (Extended Data Fig. 4), enabling the simultaneous capture of local signal features and long-range dependencies. Input features comprise 200 raw signals (+-100 centered on the target base) along with their corresponding 9-mer sequence context. Even with limited training epochs, OpenBase demonstrated exceptional detection precision and achieved high performance matrix across multiple datasets (Fig. 2a-b and Extended Data Fig. 5a-b).

**Fig. 2:**
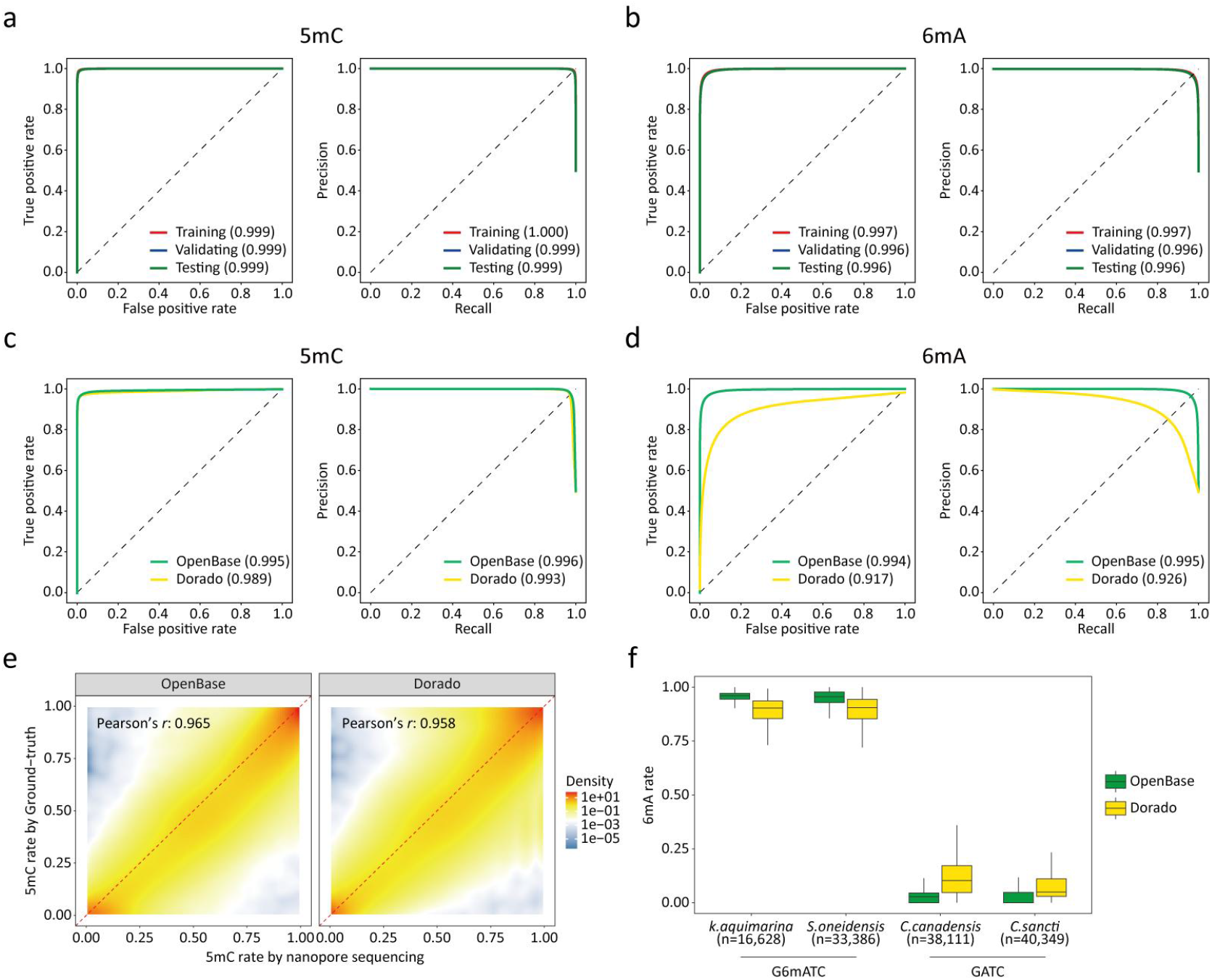
Accurate and robust single-molecule performance of OpenBase. **a-b**, Receiver operating characteristic (ROC) and precision recall (PR) curves for single-molecule 5mC (**a**) and 6mA (**b**) predictions across multiple datasets, with corresponding area under curve (AUC) values indicated. **c-d**, Single-molecule performance comparison between OpenBase and Dorado for 5mC (**c**) and 6mA (**d**) evaluated using ONT’s standard datasets. **e**, Correlation of OpenBase- and Dorado-predicted 5mC rates with BS-seq ground-truth. **f**, Distribution of 6mA rates for GATC motifs across different bacterial species as predicted by OpenBase and Dorado.

We subsequently conducted a systematic benchmarking using publicly available third-party datasets. On the standard single-molecule site-specific modification datasets produced by ONT (Oxford Nanopore Technology)^13^, OpenBase outperformed the official caller Dorado, exhibiting superior detection performance for both modifications, with a particularly marked improvement in 6mA (Fig. 2c-d). In the real biological sample HG002 (a human reference cell line from the Genome in a Bottle consortium), the predicted 5mC methylation rate at individual sites by OpenBase were highly consistent with BS-seq data^13^ and showed higher correlation than those from Dorado (Fig. 2e), confirming its reliability in complex genomic contexts. In bacterial restriction-associated 6mA detection^14^, OpenBase accurately recapitulated strain-specific methylation patterns: bacteria harboring the modified motif GATC exhibited near-complete methylation the expected sites, whereas non-modified strains displayed virtually no methylation signal (Fig. 2f). Collectively, these results demonstrate that OpenBase surpasses existing industry standards to enable high-accuracy detection of multiple non-canonical bases at the single-molecule level.

Because deep learning model performance strongly depends on data scale, we next evaluated the impact of training dataset size on OpenBase’s performance. Model improved modestly with small datasets but increased sharply once the data volume exceeded a critical threshold, beyond which improvements plateaued (Extended Data Fig. 5c). Interestingly, the inflection points differed across modification types: 6mA (∼500,000 samples) required more training data than 5mC (∼100,000 samples). This likely reflects the broader influence of 6mA on nanopore current signals, necessitating a larger dataset to capture sufficient sequence diversity.

We further extended OpenBase to train and evaluate detection models for a wide range of non-canonical bases, including selected epigenetic modifications, damage-derived bases and synthetic analogs. These non-canonical bases differ from their canonical counterparts in multiple ways: some carry additional chemical groups (e. g., 5mC and 5hmC), others harbor group substitutions that alter atomic composition (e. g., BrdU), and still others lack specific moieties (e. g., abasic site). Remarkably, OpenBase consistently achieved excellent performance metrics across all these chemically distinct categories (Fig. 3a). At the default probability threshold of 0.5, the confusion matrices clearly demonstrate that OpenBase attains high specificity and sensitivity for every type of non-canonical base tested (Fig. 3b). Notably, OpenBase exhibited exceptional resolution in discriminating between closely related cytosine derivatives (Fig. 3c). Despite the subtle signal differences between 5mC, 5hmC, and dU, the model accurately deconvoluted these variants with minimal cross-talk at the single-molecule level. This robust performance, irrespective of whether the base gains, loses, or replaces chemical groups, underscores the strong generalization capacity and broad applicability of OpenBase as a universal framework for detecting diverse non-canonical bases at the single-molecule level.

**Fig. 3:**
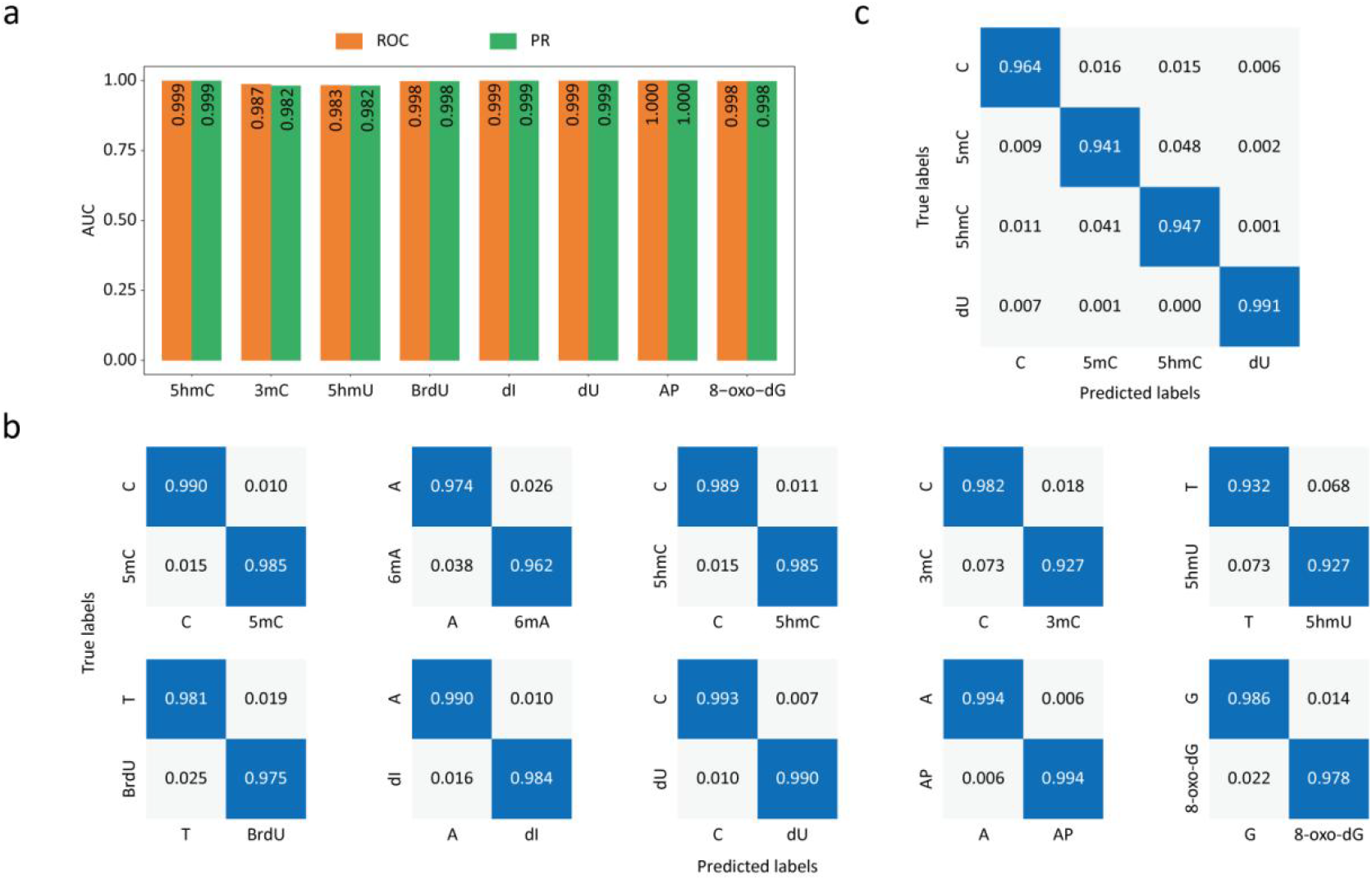
Universality of OpenBase across diverse classes of non-canonical bases. **a**, AUC values for single-molecule predictions of eight additional non-canonical bases in testing datasets. **b**, Confusion matrices of single-molecule predictions for ten non-canonical bases in testing datasets at a threshold of 0.5. **c**, Confusion matrix of single-molecule predictions among cytosine-related non-canonical bases.

In summary, OpenBase establishes an open and universal framework for single-molecule detection of non-canonical DNA bases. By integrating standardized data generation with deep learning-based modeling, it enables precise and generalizable recognition across DNA bases with diverse chemical properties, including endogenous modifications, damage-derived bases and synthetic analogs. Although this study used data from the ONT platform, the underlying design of OpenBase is platform-agnostic and can be adapted to other nanopore sequencing systems that capture base-dependent electrical signals. Its modular structure allows any base type to be profiled by simple substitution in the synthetic core region, providing a scalable foundation for exploring uncharacterized bases. Looking ahead, OpenBase may be extended to identify non-canonical bases in RNA and to achieve simultaneous detection of multiple base types. Therefore, OpenBase provides an openly accessible and generalizable platform for epigenetic analysis and synthetic biology, offering broad potential to advance single-molecule nucleic acid research^15-19^.

## Methods

### Data collection

The training datasets were generated by this study, while the evaluation datasets were derived from publicly available sources:

1. ONT’s standard datasets. Nanopore sequencing data, reference sequence and site-specific modification results were obtained from Oxford Nanopore Open Data Registry^13^.
2. HG002. Nanopore sequencing data and BS-seq results were both obtained from Oxford Nanopore Open Data Registry^13^.
3. Bacteria. Nanopore sequencing data and reference genome sequence were obtained from an AWS storage bucket^14^.

#### *In-vitro* synthesis and duplex formation of oligonucleotides

Custom oligonucleotides bearing 5’ phosphorylation were synthesized by a commercial supplier and purified via high-performance liquid chromatography (HPLC). Equimolar amounts of each complementary pair were annealed and extended using the Taq Hot Start DNA Polymerase kit (NEB, #M0495). Thermal cycling consisted of an initial denaturation at 95 °C for 30 s, followed by simultaneous annealing and extension at 68 °C for 6 min, and a final hold at 4 °C. The resulting duplex DNA products were purified with the DNA Clean & Concentrator-5 kit (Zymo Research, #D4014) according to the manufacturer’s instructions, eluted in nuclease-free water, and quantified with a Qubit fluorometer for downstream nanopore library construction.

### Nanopore sequencing and data processing

Sequencing was conducted following manufacturer’s instructions provided by Oxford Nanopore Technologies (Oxford, UK) using SQK-LSK114 library preparation kit and PromethION flowcells (FLO-PRO114M), except that the DNA repair and end-preparation steps were omitted, and adapter ligation and purification were carried out directly. Raw signal files in pod5 format were basecalled with Dorado (v0.9.0) using the parameter “--emit-moves”, generating BAM files. These Bam files were then converted to fastq format with Samtools (v1.19.2) using the command “fastq -T “*”“ to retain all tags. The resulting reads were aligned to the reference fasta using Minimap2 (v2.28-r1209) with the parameters “-ax map-ont -k 10 -w 6 --score-N 0 -y --MD --secondary=no”. Finally, Samtools was used to filter unmapped reads and supplementary alignments, convert sam files to bam format, and perform sorting.

### Feature extraction

Remora^20^ (v3.3.0) dataset prepare module was used to refine the signal mapping results with the parameters “--refine-kmer-level-table 9mer_levels_v1.txt --refine-rough-rescale --refine-scale-iters 5 --kmer-context-bases 6 2 --focus-reference-positions position.bed”, The file “9mer_levels_v1.txt” was obtained from the official ONT release, and “position.bed” specifies the locations of the target bases on the reference sequence. Based on the output generated by Remora, we extracted the raw signals corresponding to the target bases and their flanking regions, as well as the associated 9-mer sequence contexts.

### Model architecture

The raw signals and their corresponding One-Hot encoded sequences were concatenated along the second dimension and fed into a five-layer one-dimensional convolutional neural network (CNN) with channel sizes of (64, 64, 128, 192, 256) and kernel sizes of (1, 1, 2, 1, 2). The resulting feature maps were appropriately transposed and then passed through a three-layer Transformer encoder with eight multi-head attention modules. After another transposition, the outputs were processed by global average pooling along the second dimension and subsequently passed through a fully connected layer with 256 and 128 neurons. Following a dropout layer (rate = 0.3), the features were further fed into two fully connected layers with 128, 64, and 2 neurons, respectively. The final output probabilities were obtained through a Softmax activation function.

### Training and evaluation

#### Training

Datasets corresponding to non-canonical bases and their respective canonical counterparts were merged, balanced, and randomly divided into training, validation, and testing sets at a ratio of 9:0.5:0.5. The models were implemented in PyTorch and trained using the Adam optimizer (lr=1e-4, weight_decay=0.01, and betas=(0.9, 0.999))to minimize the cross-entropy loss. A batch size of 512 was used, and training was performed for 20 epochs. After each epoch, model performance was evaluated on different datasets using metrics including ROCAUC, accuracy, sensitivity, and specificity.

#### Evaluation

For the standard datasets provided by ONT, both the 5mC and 6mA datasets contain corresponding modifications introduced at fixed positions, while the control dataset contains no modifications. This allows us to plot ROC and PR curves and calculate AUC values based on single-molecule predictions at specific sites across different datasets. For the HG002 dataset, we compared the methylation rates derived from single-molecule 5mC predictions with the BS-seq ground truth and calculated the Pearson correlation coefficient (r), using sites on chromosome 1 with a minimum coverage of 20. For the bacterial datasets, we computed the methylation rates of all A sites within the GATC motif across four bacterial species, based on the single-molecule 6mA predictions. To establish a comparative baseline, Dorado (v0.9.0) was employed for basecalling with specified modification models. Modification states were subsequently parsed using modkit (v0.3.1); specifically, the pileup and extract modules were utilized to derive site-level methylation rates and single-molecule modification status, respectively.

## Data availability

The sequencing data generated in this study have been deposited in the European Nucleotide Archive under accession number PRJEB100817. ONT’s standard datasets, as well as the ONT dataset and BS-seq results of HG002, are available from the Oxford Nanopore Open Data Registry at the following AWS storage buckets: s3://ont-open-data/modbase-validation_2024.10, s3://ont-open-data/giab_2023.05/flowcells/hg002/20230424_1302_3H_PAO89685_2264 ba8c, and s3://ont-open-data/gm24385_mod_2021.09/bisulphite, respectively. The ONT datasets and reference genome sequences of the four bacterial species were obtained from s3://cultivarium-publication-data/MICROBEMOD-DATA-NOV2023.

## Code availability

The source code and detailed tutorial for OpenBase are available on GitHub: https://github.com/zhongzhd/OpenBase.

---

**Extended Data Fig. 1:**
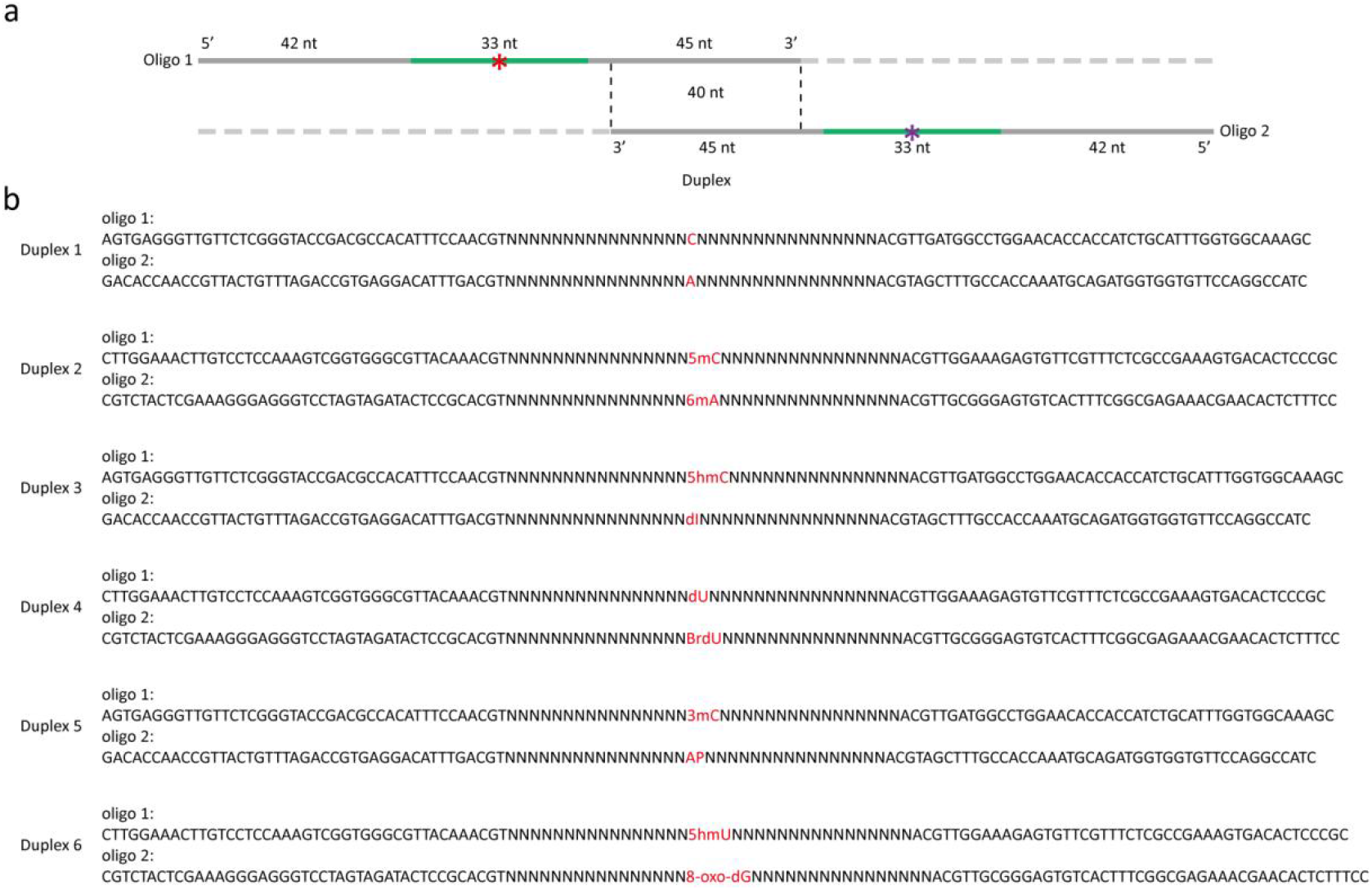
Design of standard data. **a**, Lengths of different regions of the single-stranded oligonucleotide. **b**, Template sequences corresponding to different bases.

**Extended Data Fig. 2:**
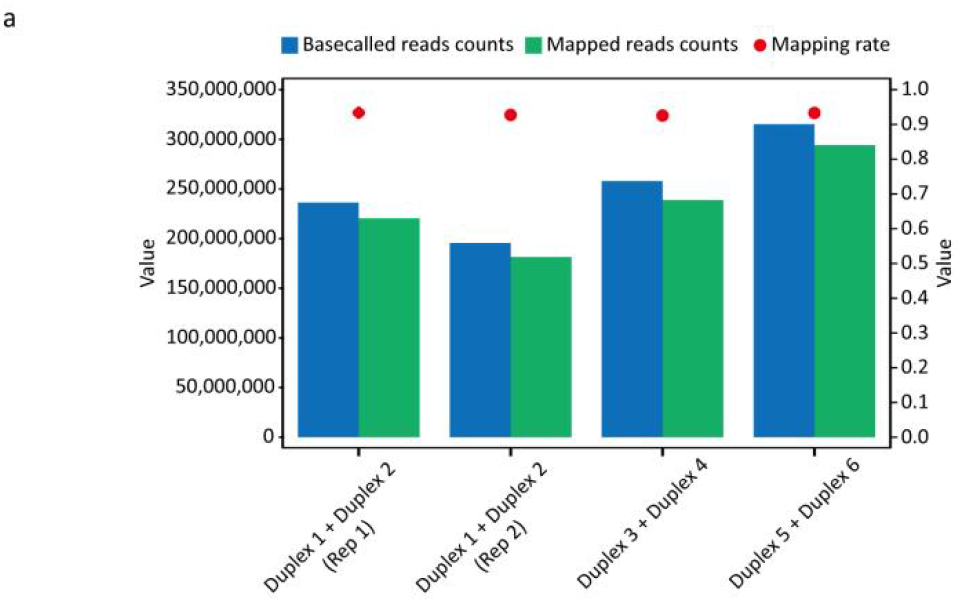
Alignment statistics for different sequencing libraries.

**Extended Data Fig. 3:**
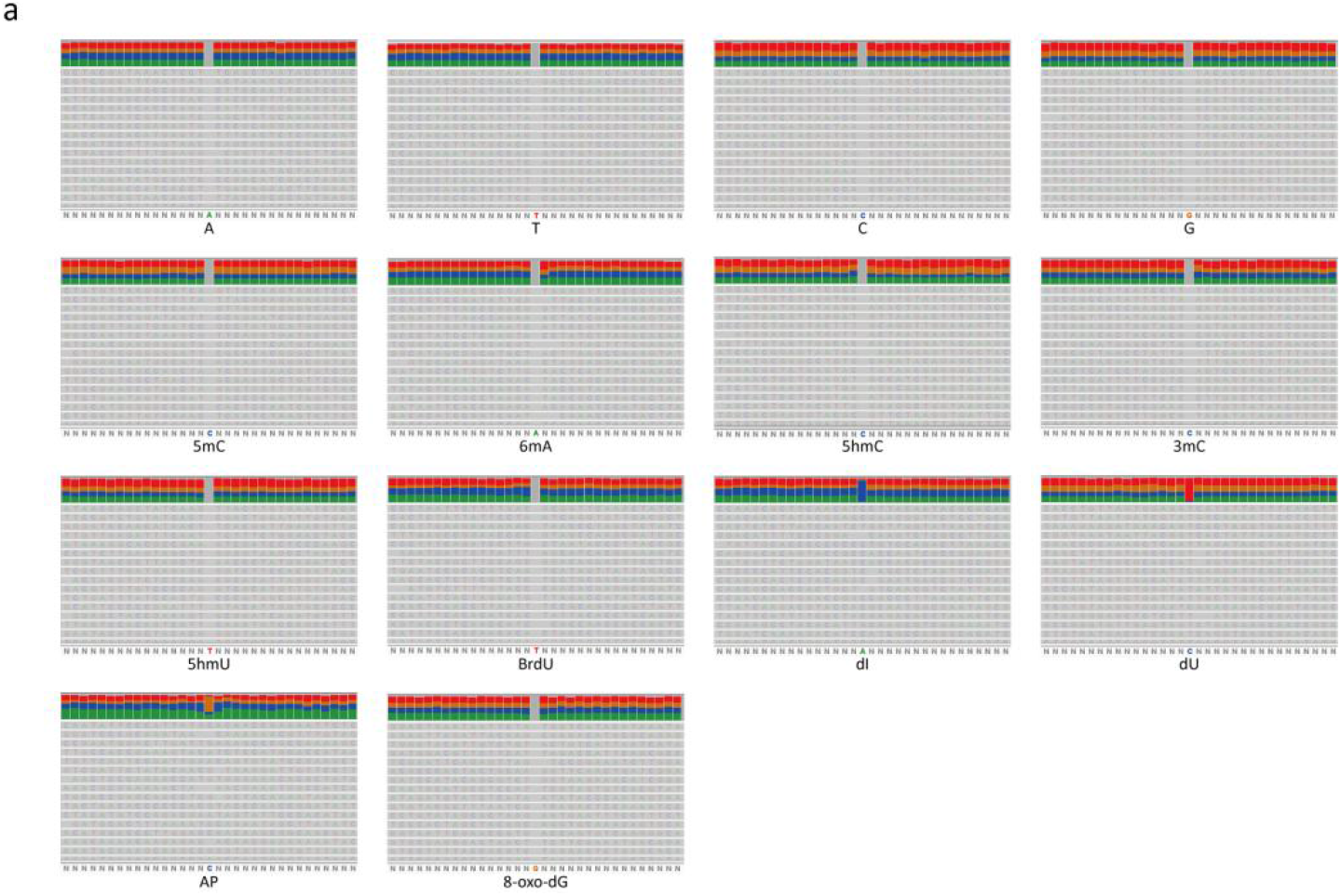
IGV snapshots illustrating the diversity and uniformity of random regions in the synthetic sequences for 14 base types.

**Extended Data Fig. 4:**
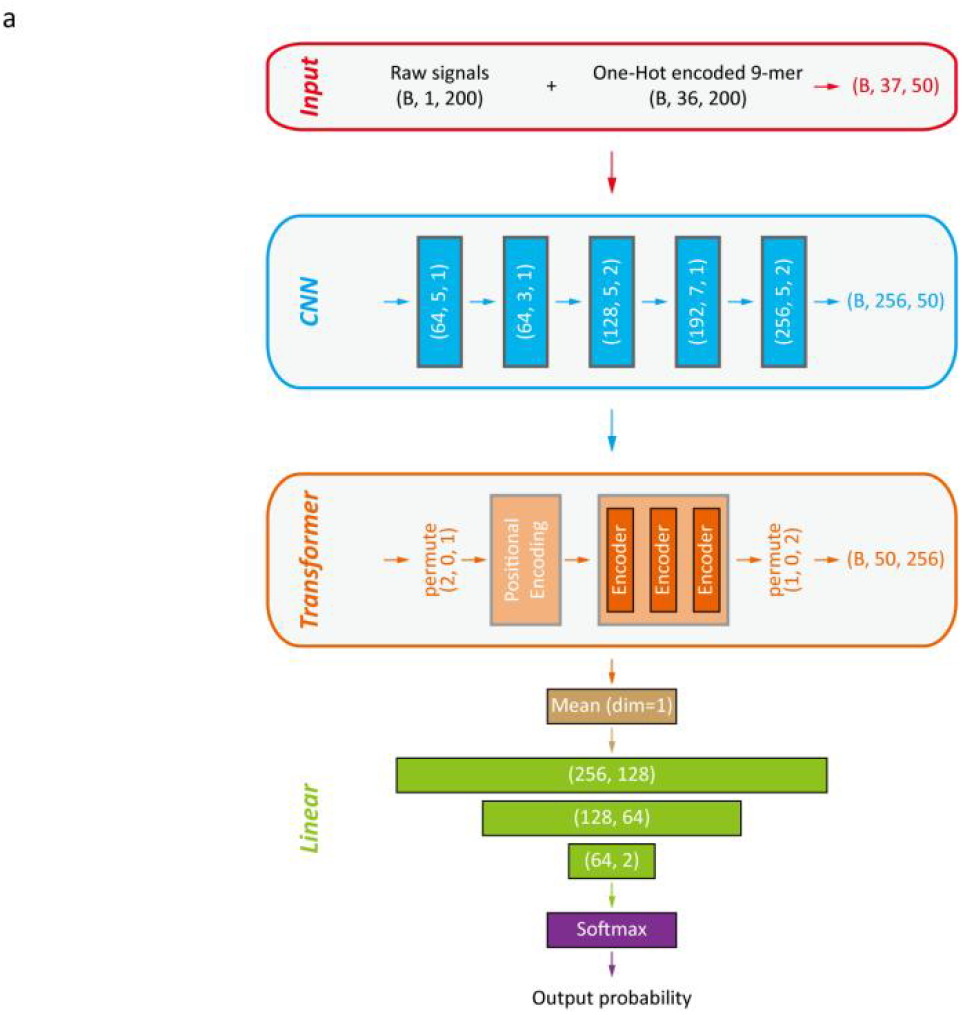
Schematic of the deep learning model architecture used in OpenBase.

**Extended Data Fig. 5:**
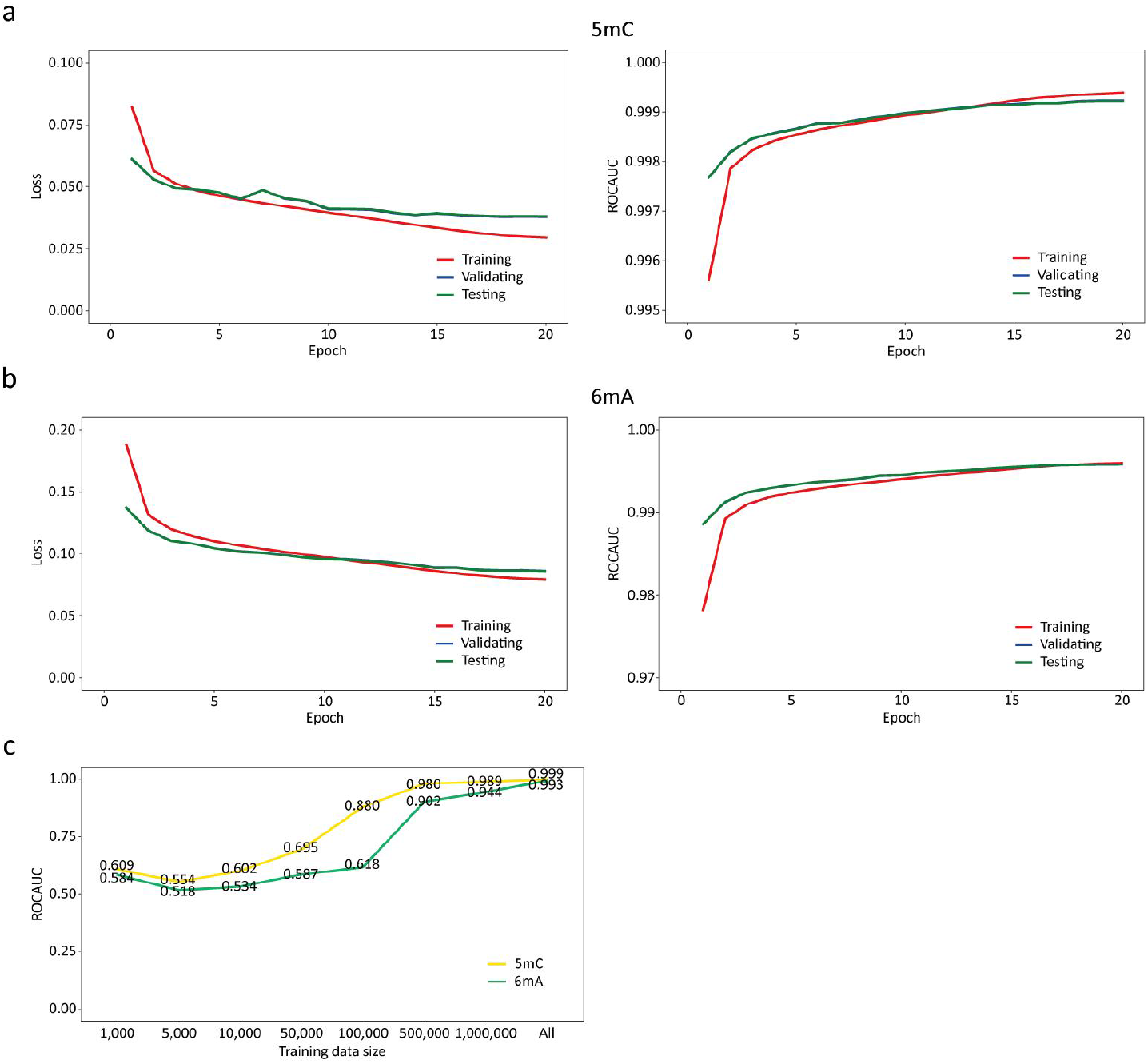
Training dynamics and data scalability of OpenBase. **a-b**, Variation of loss and ROCAUC with training epochs for OpenBase predictions of 5mC (**a**) and 6mA (**b**) across multiple datasets. **c**, ROCAUC values for 5mC and 6mA predictions under varing training data sizes.

